# Memory of split-belt walking endures weeks in young children and younger adults but less so in older adults

**DOI:** 10.1101/2021.02.12.430812

**Authors:** Brittany Lissinna, Allison Smith, Kaylie La, Jaynie F Yang

## Abstract

Adults and children modify how they move to accommodate persistent changes in their surroundings, called motor adaptation. Walking-related motor adaption can be seen when one walks on a treadmill with two belts running at different speeds, a split-belt treadmill, where a new walking pattern is slowly adopted with practice. This has been suggested as a way to improve left-right symmetry in walking after a stroke. Central to using split-belt walking for rehabilitation is whether the adapted motor pattern is retained over days and weeks, and whether this motor memory is a function of the person’s age. When first exposed to split-belt walking, the walking is asymmetric (initial error), resembling a limp. With subsequent exposure, the error is smaller than the initial error, indicating the adaptation was remembered. Here, we explored the persistence of this memory over 24 hours, one week, and two weeks, in young children (3-6 yr old), younger adults (20-30 yr old) and older adults (50-70 yr old). We found that the motor memory declines with the interval duration, but was still evident up to 2 weeks after initial exposure. Retention of the motor memory in children and younger adults was better than older adults. Further, forgetting between trials was seen on the first day of split-belt walking especially in children and older adults, but this forgetting diminished with repeated trials. The results indicate that long-term memory of motor adaptation on the split-belt treadmill is affected by age, but it may be possible to enhance the memory by more frequent and repeated exposure. This suggests that when using split-belt adaptation in rehabilitation, the sessions should ideally be less than one week apart.

## 1. Introduction

Motor adaptation is a form of motor learning that is characterized by the modification of a well-learned movement. An example is dart throwing to a target while wearing prism glasses that shift the visual image in one direction. The throwing movement is initially inaccurate (initial error), but is gradually modified over repeated trials, called motor adaptation (Martin et al., 1996). When the condition is returned to normal by removing the prism glasses, a movement error opposite to the initial error is seen, called aftereffect. The aftereffect indicates that the motor pattern had been modified. Motor adaptation may be a potential intervention for certain types of movement problems, such as using prism glasses to reduce hemi-neglect after stroke (Mizuno et al., 2011), or split-belt walking to correct walking asymmetry after stroke (Betschart et al., 2018; Reisman et al., 2013).

We studied split-belt walking, which is walking on a treadmill with a belt for each leg, running at different speeds. When first exposed to split-belt walking, the person appears to be limping because a large asymmetry is induced in the left and right steps (Reisman et al., 2005). Early in adaptation, the leg on the fast belt is pulled further back than the leg on the slow belt, creating a spatial and timing difference between the left and right steps (Bruijn et al., 2012; Reisman et al., 2005). With continued walking, the fast leg learns to reach forwards further and faster, equalizing the two steps. These changes result from modifications to the activation of leg muscles bilaterally (Vervoort et al., 2020). Retention of this learning is shown by a smaller asymmetry in walking when participants were re-exposed to the same task after a period of rest (Roemmich & Bastian, 2015). Further, returning the belts to the same speed (i.e., tied-belt speeds) after adaptation results in the opposite asymmetry seen during initial exposure to the split-belt, an aftereffect (Reisman et al., 2005). Split-belt adaptation may correct walking asymmetry after adult stroke (Betschart et al., 2018; Reisman et al., 2013), and it is evident at a very young age (Musselman et al., 2011; Vasudevan et al., 2011), suggesting that young children who suffer a stroke around birth (Dunbar & Kirton, 2018) may also benefit from this intervention.

One aspect of motor adaptation that is less well understood, and indeed central to using motor adaptation as an intervention, is how well the adapted movement pattern is retained over longer periods of time, such as days and weeks. Thus, as an initial step, we explored how well split-belt adaptation is retained by people of different ages with intact nervous systems. We were especially interested in studying both very young children and older adults, since both groups may show differences in the retention of motor adaptation compared to younger adults. For example, typically developing young children show slower motor adaptation during the first few hundred steps of split-belt walking compared to adults (Musselman et al., 2011; Vasudevan et al., 2011). In some other forms of motor adaptation, a slower adaptation rate has been hypothesized to be better retained than faster adaptation rate (Joiner & Smith, 2008). Older adults, however, show poorer retention within a session when short breaks are interspersed (Malone & Bastian, 2016), which may suggest poorer retention over longer periods. This is an exploratory study on the retention of split-belt adaptation over days and weeks.

## 2. Methods

### 2.1 Participants

Participants with no known health problems that would interfere with motor learning were recruited. We further confirmed they were naïve to split-belt walking. We chose to compare three age ranges: 3-6 years old, because they showed adaptation beginning around this age (Vasudevan et al., 2011), 20-30 years olds because they have been best studied, see review (Torres-Oviedo et al., 2011), and >50 year olds because of their poorer retention within a session (Malone & Bastian, 2016). The small age range for young children was to minimize the variability associated with development (Vasudevan et al., 2011). The older age range is associated with higher risk of pathologies with walking asymmetries, such as from stroke. Thus, the information will provide a comparison for the uninjured, older population.

Adult participants and a parent/guardian of children gave informed, written consent. Verbal assent was obtained from the children at the time of the experiments. The Health Research Ethics Board, University of Alberta, approved the study (Pro00043622). There was no prior information regarding possible differences between groups. Thus, sample size was based on a prior publication of age-related differences in split-belt walking (Vasudevan et al., 2011), which was approximately 10 participants per group, in line with our group sizes (see Table 1 in Results).

**Table 1.** Participant numbers and ages from each experimental groups.

| Gap duration |  | 24 hours | 1 week | 2 weeks |
| --- | --- | --- | --- | --- |
| Kids | N (f) | 12(6) | 9(3) | 9(3) |
|  | Age (yr) | 4.6±0.9 | 4.9±0.9 | 4.4±0.5 |
| Young adults | N (f) | 7(4) | 7(2) | 7(4) |
|  | Age (yr) | 23.5±3.7 | 24.0±2.0 | 25.6±2.3 |
| Old adults | N (f) | 7(4) | 8(6) | 7(4) |
|  | Age (yr) | 62.1±8.6 | 59.7±5.7 | 59.7±7.1 |

N is the number of participants contributing data in the group with the number of female (f) participants in parentheses. Age is represented by years (yr) with mean ± 1 SD.

### 2.2 Experimental Protocol

Testing was performed on 2 days, separated by 24 hours, 1 week or 2 weeks (Figure 1). These intervals were chosen because retention after 24 hours is known for children and adults (Day et al., 2018; Musselman et al., 2016) and can provide a comparison with our findings. One and two weeks were chosen because these intervals are realistic session intervals for those interested in using motor adaptation in rehabilitation. The allocation of participants to the gap intervals was partly determined by participant availability, but was otherwise random.

**Figure 1.**
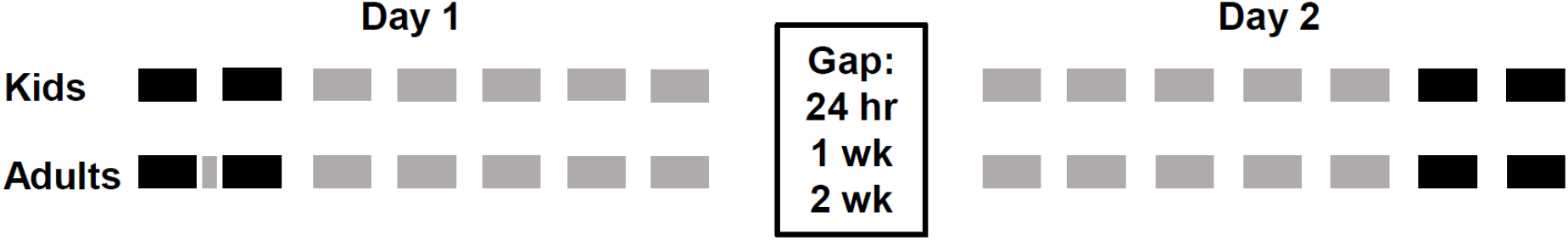
Experimental sequence. Each walking trial is represented by the solid horizontal bars. Black bars represent tied-belt trials and gray represent split-belt trials. All trials were 3 min long except the surprise trial of 5 strides, 5 seconds long, short gray bar between 2 tied-belt trials on Day 1. All participants experienced the same sequence of trials, with the exception of the surprise trial for adults only. Three gap durations between Day 1 and Day 2 were used for each of the age groups.

On Day 1, two tied-belt trials (belts at the same speed) were followed by 5 split-belt trials (speed ratio 2:1) (Figure 1). All trials were 3 min long, separated by a 1 min standing break. The trial length was to accommodate children, who were more amenable to multiple short bouts rather than continuous walking. Because some adults exhibit surprise when first exposed to split-belt walking, causing an exaggerated initial error, a ‘surprise’ trial (≤5 s) was inserted between the two tied-belt trials as is common in other studies (Malone et al., 2011). Children do not display ‘surprise’ (Musselman et al., 2011; Vasudevan et al., 2011), so a surprise trial was not used for children. On Day 2, participants started with 5 split-belt trials to determine their retention of the adaptation, followed by 2 tied-belt trials to quantify the aftereffect.

A split-belt treadmill (Woodway USA Inc., Waukesha, WI) with the belts controlled by separate motors was used. Because of the enormous difference in leg length between children and adults and among children, the speed of the tied-belt trials was set according to leg length for children, with the distance from the greater trochanter to the lateral malleolus (m) providing the speed in m/s, as previously reported (Vasudevan et al., 2011). The speed for adults was set at a tied-belt speed of 0.7m/s, and split-belt speeds of 1.4 m/s and 0.7 m/s. The slow speed corresponds roughly to the leg length of an average adult woman, i.e., approximately 0.7 m, thereby making it similar to how the slow speeds were determined for children. The particular speed combination has been used in a number of split-belt studies (Vasudevan & Bastian, 2010)(Malone et al., 2011)(Roemmich & Bastian, 2015), which will also facilitate comparisons. The leg on the fast belt was chosen randomly on the first day, and maintained for Day 2.

For safety, all participants held a front handrail and an adult spotter stood behind the children. All participants watched a movie. Adults were instructed not to think about their walking and to focus on the movie. Children were not always focused on the movie, so if we noticed they were watching their feet and playing, we used questions about the movie to divert their attention from walking.

### 2.3 Instrumentation

Infrared emitting markers (Northern Digital Inc., Waterloo, ON) were positioned bilaterally at the top of the pelvis directly above the hip, the greater trochanter, the knee joint line, the lateral malleolus, and the head of the 5^th^ metatarsal. Two 3-D Investigator motion sensors on each side of the treadmill captured the positions of the markers using NDI software. In addition, a video camera, synchronized in time with the marker data, allowed the exclusion of steps when participants were not stepping with one foot on each belt.

### 2.4 Data Analysis

We focused on step length symmetry, because it is a global measure of spatial and temporal symmetry on the split-belt (Vasudevan et al., 2011). Custom MatLab (The Mathworks Inc., Natick, MA) code was used to analyze the data. Step length was defined as the antero-posterior distance between the ankle markers of each leg at the time of heel strike of the slow (SLS) and fast (SLf) leg, respectively. Step symmetry (SS) was the normalized difference between the step lengths:

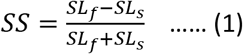

A negative SS means the slow step was longer than the fast step. Individual participant data were smoothed using epochs of 9-step averages, arbitrarily chosen as a compromise between the fast and slow time courses of learning in adults and children, respectively. The data were then averaged across participants within each age group and time gap.

The differences in the time course of adaptation on Day 1 between the age groups were determined by collapsing the data across the 3 gap durations for each age group. This was justified since there were no differences in age between the participants with different gap periods, and Day 1 was collected in exactly the same way for all participants.

Step symmetry during specific times were of interest. Baseline symmetry was the average from all steps in the second tied-belt trial on Day 1. The initial error on the split-belt for Day 1 (IE_1_) and Day 2 (IE_2_) were the average of the first 9 steps on the split-belt each day. The final error on Day 1 (FE_1_) was the average of the last 30 steps on the split-belt, in line with prior studies (Malone et al., 2011) to provide a stable estimate for final plateau in adaptation. Forgetting between Day 1 to Day 2 was the difference between the initial error on Day 2 and the final error on Day 1, with negative values indicating forgetting (Forgetting = IE_2_-FE_1_). The aftereffect (AE2) was the average of the first 9 steps of the tied-belt on Day 2.

Since young children do not always show adaptation (Vasudevan et al., 2011), we first sought evidence of adaptation in each child by determining if the average step symmetry of the last split-belt trial was significantly more positive than the first split-belt trial on Day 1 (t-test, p<0.05). Further, presence of an aftereffect on Day 2 was confirmed by the average step symmetry of the first 9 steps on the tied-belt being more positive than the last 30 steps on the split-belt (t-test). If both of the above comparisons showed no differences, indicating absence of learning, the child’s data was removed from further analysis. This was justified, because including children who were not adapting would mean that they would show perfect retention (i.e., Forgetting equals zero), when in fact they did not adapt at all. This would unfairly bias the results towards better learning in children, which would unfairly support our prediction.

Forgetting between trials on Day 1 was the difference between the average of the 9 steps just before and just after a break. Since Day 1 was the same for all gap duration, the data were collapsed across gap durations for each age group.

### 2.5 Statistical Analysis

One-way ANOVAs were used to compare the ages between participant groups within the same gap interval. Because there were no age differences within each age group, the time course of adaption on Day 1 was collapsed across different gap durations within each age group. This data were then modeled by a mixed, linear regression model to compare participants in the different age groups using the log likelihood ratio test. In addition, we used the Akaike Information Criteria to determine if a linear or exponential model provided a better fit of the time course for each age group.

Two-way ANOVAs (factors: age group and gap interval) determined the differences between the groups in: 1) baseline stepping symmetry, 2) initial error on the split-belt, 3) aftereffect in Day 2, and 4) forgetting between Day 1 and Day 2. Within-day forgetting for Day 1 was compared between the age groups using a mixed model repeated-measures ANOVA (within-subject factor: breaks, between-subject factor: age group). Because this is an initial exploratory study, we wished to minimize the chance of Type II errors. Thus, a significant difference was set at p<0.1 for all comparisons

## 3. Results

### 3.1 Participants

Seventy-four volunteers (31 children, 21 younger adults, 22 older adults) participated in the study. Data from one child were eliminated from analysis because of an absence of motor adaption. See Table 1 for participant numbers, sex and age in each group. All participants completed the trials except for one child, who did not want to continue after 4 split-belt trials on Day 1; this same child completed all trials on Day 2. This child’s data were included for all analyses except for the forgetting within Day 1, in which there was missing data for the 4^th^ break (i.e., no trial #5).

The age of the participants within each age group were the same (Table 2, Age). Symmetry of walking with tied-belts on Day 1 was the same for all 9 groups (Table 2, Baseline step symmetry), as was the initial error on the split-belt on Day 1 (Table 2, SB Initial error), indicating all groups were equally perturbed by the split-belt.

**Table 2.**
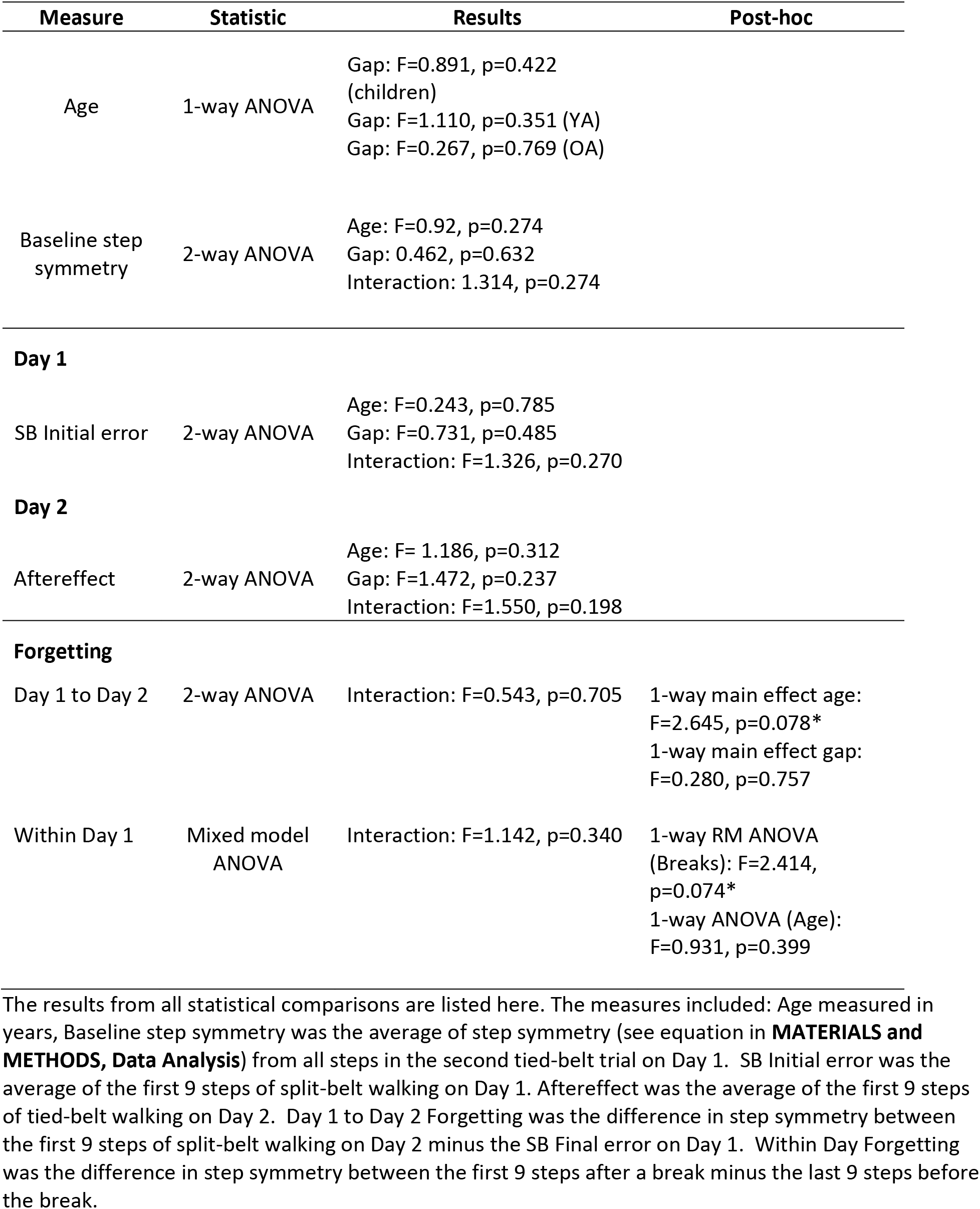
Statistical results.

### 3.2 Time course of adaptation and overall learning

The effect of age on the time course of adaptation during split-belt walking on Day 1 is shown in Figure 2. The time course is truncated at the lowest number of total steps obtained. All groups showed a reduction in step asymmetry over time, as expected (Reisman et al., 2005), with the children showing a slower time course and more variability compared to the adults, confirming previous findings (Vasudevan et al., 2011). The best fitting model for all age groups was a linear model. The age groups were significantly different from each other, with children being different from younger adults [log ratio(LR) chi2 (62)=462.65, prob=0.0000] and older adults [LR chi2(62)=180.36, prob=0.0000], but younger and older adults were not different from each other [LR Chi2(62)=-282.29, prob=1.000]. The aftereffect on Day 2, a reflection of overall learning over the two days, was the same across all groups (Table 2, Day 2, Aftereffect), indicating a similar amount of learning for all age groups.

**Figure 2.**
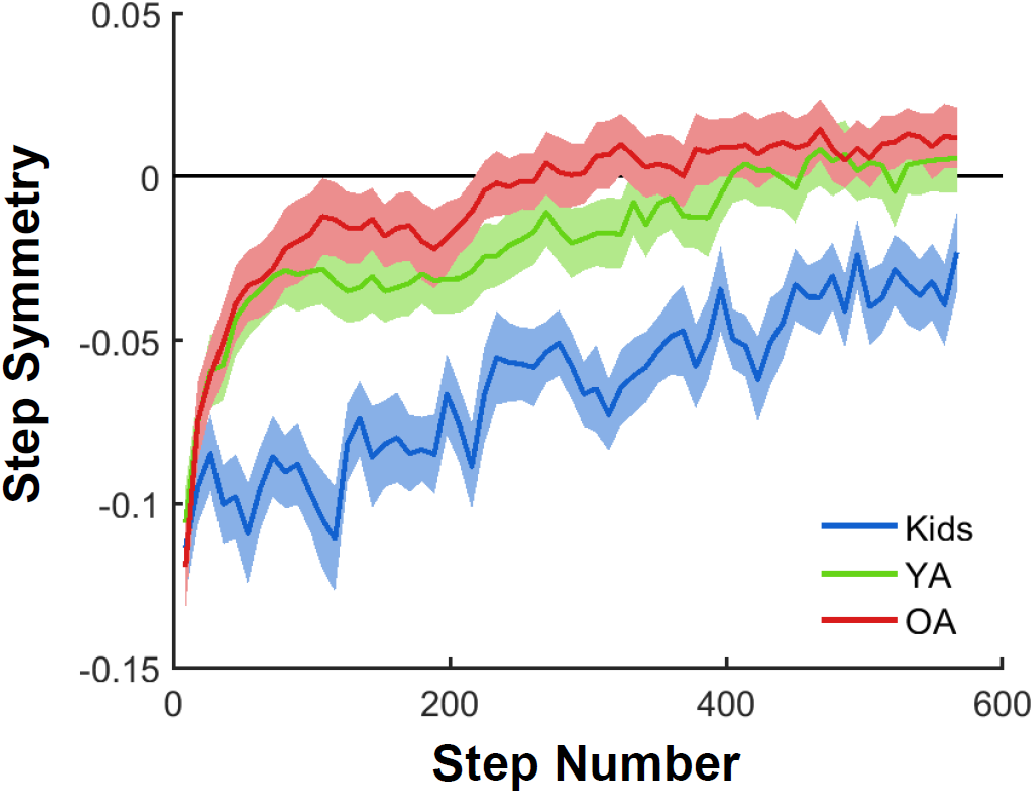
Time course of split-belt adaptation on Day 1. Average (9-step average) step symmetry is shown for the 3 age groups over the adaptation period of ~15 min. The age groups each contain data from n=30 for young children (Kids), n=21 for younger adults (YA), and n=22 for older adults (OA).

### 3.3 Forgetting between Day 1 and Day 2

The time course of split-belt adaptation is superimposed for each of the groups on Day 1 and Day 2 in Figure 3. All groups showed evidence of retention on Day 2, as reflected by the starting steps on Day 2 (orange) being less asymmetric than the starting steps on Day 1 (blue). The smaller initial error on Day 2 led to all groups achieving symmetric walking sooner on Day 2 compared to Day 1. The forgetting between days is summarized in Figure 3. Average forgetting at 2 weeks was 42%, 52% and 67% for children, younger and older adults, respectively, estimated by dividing the average initial asymmetry on the split-belt on Day 2 by the average initial error on the split-belt treadmill on Day 1.

**Figure 3.**
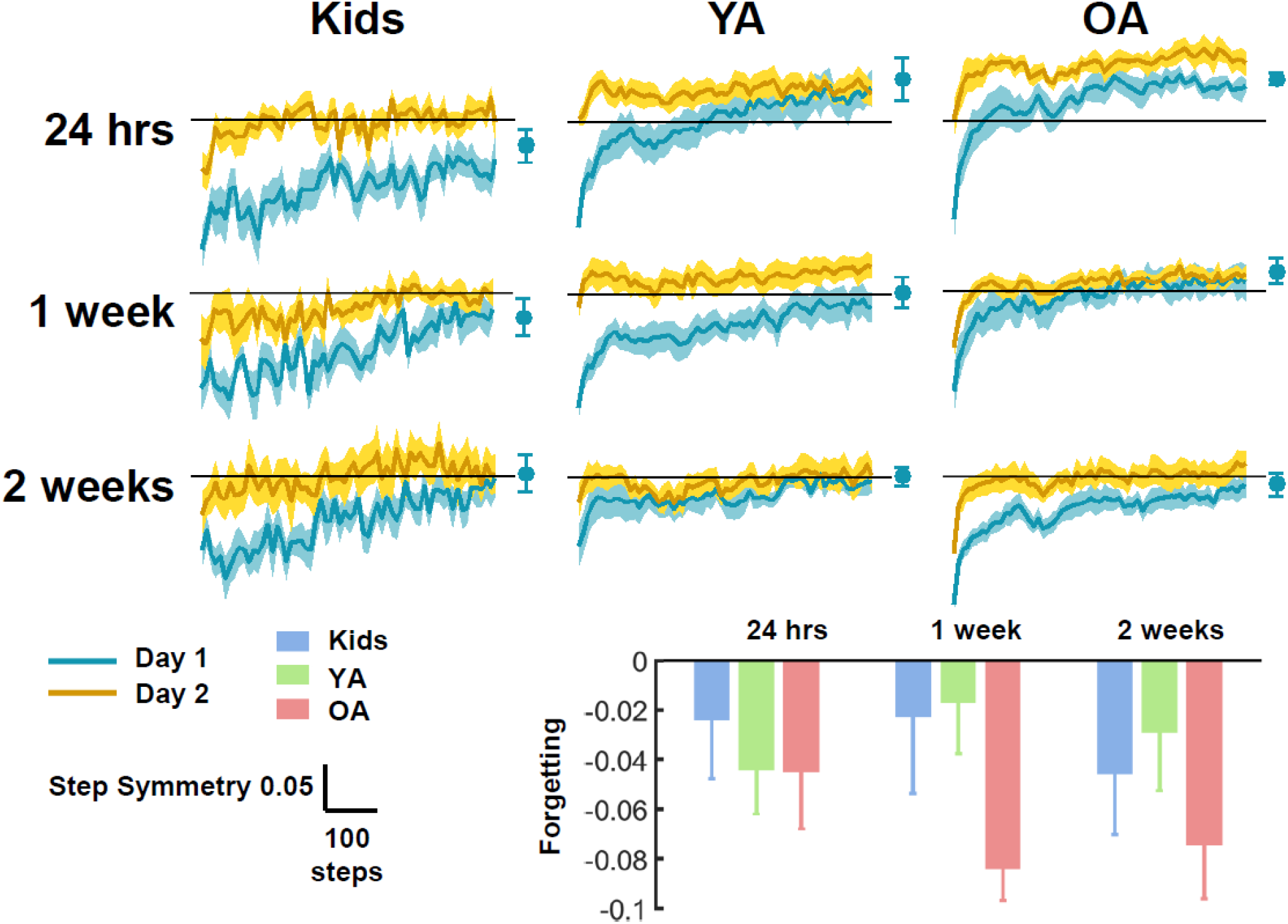
Average split-belt adaptation on Day 1 and Day 2 for each age group and gap durations. Adaptation is shown for 9 step averages across 567 steps, with the final step symmetry on Day 1 (average of last 30 steps, final error) shown as a blue symbol with ±1 SEM. The bar graph shows the forgetting between the Initial Error on Day 2 minus the Final Error on Day 1. Means and 1 SEM are shown. Older adults (OA) showed more forgetting compared to the younger age groups (Kids and younger adults – YA). Abbreviations: Kids - children 3-6 y.o.; YA - younger adults, OA - older adults.

Statistical tests of forgetting showed no age by gap interaction (Table 2, Forgetting, Day 1 to Day 2). Thus, 1-way ANOVAs were used to examine the main effect of age and gap duration. Only age was significant (Table 2, Forgetting, Day 1 to Day 2).

### 3.4 Forgetting between trials within a day

Figure 4 shows the step symmetry before and after each break, together with the initial 9 steps on the split-belt. Statistical comparison of the forgetting within Day 1 during the breaks showed a significant sphericity (Mauchly’s test, p=0.055), so the Greenhouse-Geisser correction was used. Two-way mixed-model ANOVA indicated no significant interaction, so main effects were examined. Post-hoc 1-way repeated-measures ANOVA indicated a significant effect of breaks when collapsed across age groups (Table 2, Forgetting, Within Day 1), but no age effect when collapsed across breaks (1-way ANOVA). Subsequent post-hoc test contrasting Break 1 with other breaks using paired t-tests indicated a significant difference between Break 1 and Break 3 (p=0.040), and Break 1 with Break 4 (p=0.045), but no difference between Break 1 and Break 2 (p=0.454).

**Figure 4.**
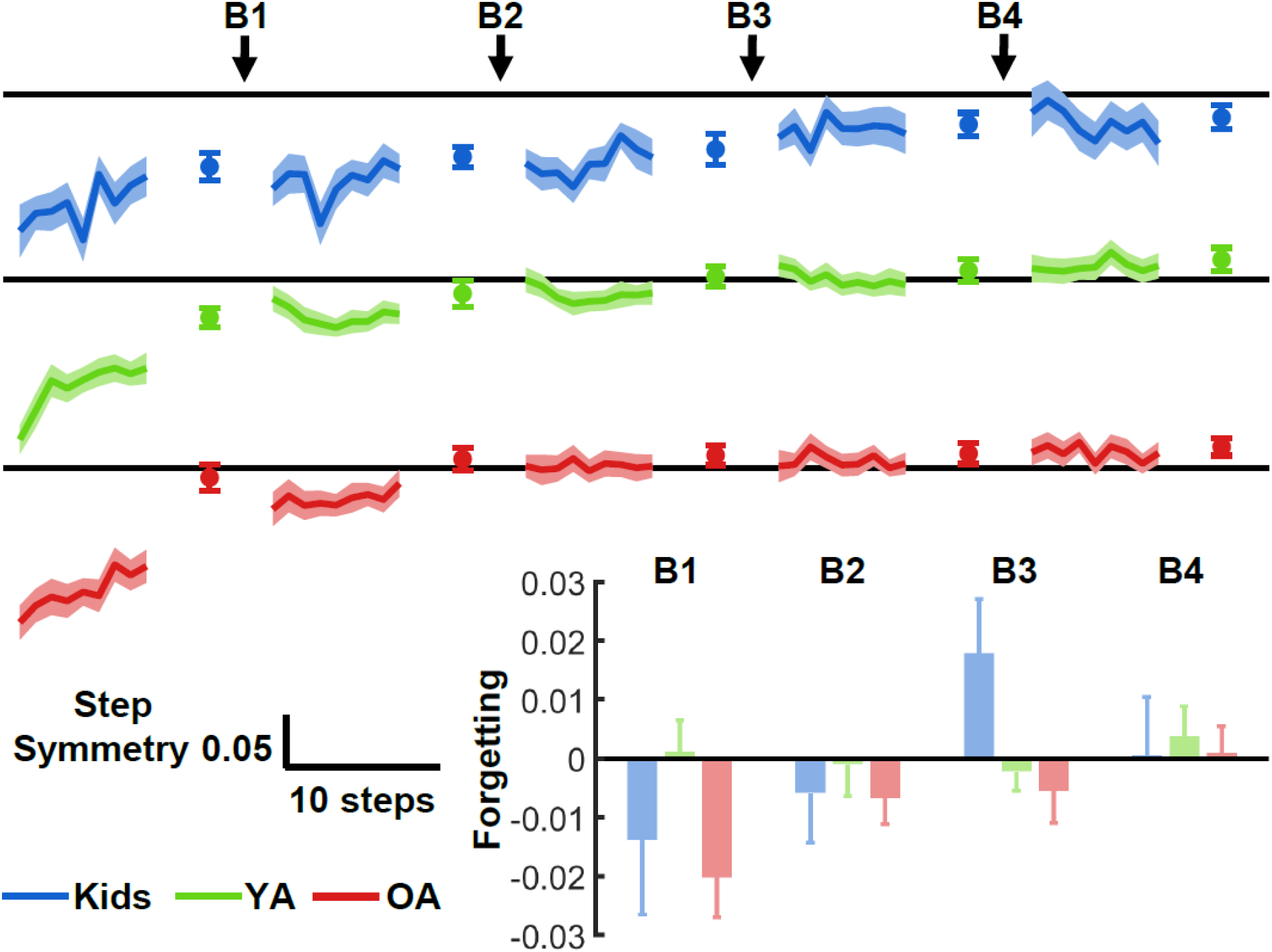
Forgetting between trials on Day 1. The step-by-step average step symmetry for each age group is shown for the first 9 steps on the split-belt treadmill, then the overall average of the last 9 steps before the breaks (B) and the step-by-step average of the first 9 steps right after the breaks, averaged across participants. The bar graph insert represents the quantified difference in forgetting between trials (forgetting = average of first 9 steps after break – average of last 9 steps before break). Kids – children, YA-younger adults, OA – older adults.

## 4. Discussion

### 4.1 Overview

The novel findings are that retention of split-belt adaptation remains strong for children and younger adults after a 1- and 2-week interval from the first exposure, whereas it was weaker for older adults. Further, within the first day of split-belt adaptation, there was a trend for children and older adults to show more forgetting at the breaks, but this forgetting was short-lived. Forgetting was similar between Breaks 1 and 2, whereas less forgetting was seen at subsequent breaks. Contrary to our original expectation, young children were no better than younger adults at retention between days, at least up to a 2-week interval after first exposure. Within a day, forgetting is reduced with repeated trials in all age groups.

### 4.2 Methodological limitations

First, we did not have any a priori information to base our sample size selection. Thus, sample size of ~10 per group (Table 1) was selected based on previous split-belt studies in which age-related differences in adaptation were addressed (Vasudevan et al., 2011). We adjusted our test of significance to be more liberal at 0.1 in this initial study. Nevertheless, the statistical power remains low, and the results should be considered preliminary.

Second, the difference in body height between young children and adults is substantial. We used leg length to attempt to equalize the experience of belt speed for children, but we did not attempt to modify the speed in the same way for adults. This is partly because the belt speeds we used for adults has been used in a variety of other split-belt studies of adults, making it easier for comparison, but also because differences in height between individual adults is proportionately much smaller than the differences in height between children and adults. In spite of this limitation, the initial error produced in all age groups was not statistically different, indicating the differential speeds resulted in similar motor errors. Thus, the starting point for adaptation is the same for all groups.

Third, both adults and children watched movies while walking, but we cannot be sure the distraction was the same between participants. Children tended to be more easily distracted compared to adults, and the novelty of the split-belt treadmill meant that some children were not always focusing on the movie, but watching their feet on the treadmill, particularly early in the adaptation. We used questions about the movie to try to draw their attention back to the movie, which was never necessary with the adults. Did asking children questions during adaptation unfairly modify their adaptation and memory compared to adults? We feel this is unlikely for a number of reasons. Distraction in younger adults does not affect the adaptation rate to split-belt walking (Mariscal et al., 2020). Distraction modified the early adaptation rate in older adults (Malone & Bastian, 2010), but did not affect the overall adaptation over the 16 min period of adaptation. Children do not show the same cognitive component of adaptation (see Section 4.3, below), characterized by the rapid phase of adaptation in the initial few minutes of adaptation. Finally, our interest was in how much adaptation was retained between days, which is a comparison of the final state of adaptation on Day 1 and the initial state of Day 2 for each group. Thus, even if the final state on Day 1 was different between groups, it should not affect the proportion of adaptation retained.

### 4.3 Characteristics of adaptation and memory in children

The rate of adaptation on Day 1 was considerably slower in children compared to adults in spite of identical initial errors, confirming previous reports for split-belt adaptation (Musselman et al., 2011; Vasudevan et al., 2011) and consistent with other forms of adaptation in children (Konczak,, J., Jansen-Osmann, P., Kalveram, 2003). The slower time course of split-belt adaptation may be related to the absence of a ‘cognitive component’ of learning, seen in adults (Huang et al., 2011; Morehead et al., 2015), since the fast time course of split-belt adaptation can be interfered with by a concurrent cognitive task, especially in older adults (Malone & Bastian, 2016). Further evidence that the cognitive component is absent in children is supported by the absence of another type of motor memory seen on the split-belt, called ‘savings’ (Musselman et al., 2016). Savings is defined as a faster relearning of the acquired motor adaptation (i.e., split-belt walking) after the initial adaptation is ‘washed out’ by an extended period of tied-belt walking (Roemmich & Bastian, 2015). Savings in motor adaptation have been hypothesized to result from a cognitive process of motor selection or retrieval (Huang et al., 2011; Morehead et al., 2015).

The slower time course of adaptation in children did not confer them with better memory, at least in our hands. We cannot discount the possibility that the higher step-to-step variability in children’s walking may have made it difficult to detect a difference, and would require more statistical power to detect. Nevertheless, this initial study provides information with which future sample sizes for these age groups can be estimated with greater statistical power.

### 4.4 Forgetting in older adults

Unexpectedly, the forgetting between trials within Day 1 was not different between the age groups, in contrast to the differences seen in a previous study, in which older adults forgot more between trials compared to younger adults (Malone & Bastian, 2016). One potential reason is the inclusion of children in our dataset, which introduced more variability within and between participants, reducing the statistical power. Also, children showed some forgetting in the first two breaks (Fig 4, bar graph inset), in contrast to younger adults, and more similar to older adults. Further procedural differences between the current study and that by Malone and Bastian include the duration of each split-belt trial, and the rest period between trials, both of which were considerably shorter in our study. Thus, instead of five 3-min split-belt trials with 1 min rest between trials here, Malone and Bastian used three 5-min trials with 5 min rest between trials. The shorter breaks may have resulted in less forgetting in this study.

### 4.5 Implications for motor adaptation as a form of therapy for walking

Split-belt adaptation has been efficacious for reducing step length asymmetry after stroke when the training was repeated over days (Betschart et al., 2018, 2020; Reisman et al., 2013). In both studies, the leg with the shorter step length is placed on the fast belt to augment the asymmetry. In one case (Reisman et al., 2013), participants were exposed to split-belt walking on average every 2.3 days for 4 weeks (12 sessions total), and in the other (Betschart et al., 2018, 2020), sessions were repeated every 2.3 – 3.5 days for 2-3 weeks (6 sessions total). These exposure frequencies are well within the range we would predict to induce good retention from all age groups our study (i.e., <1 week). Both studies used a belt speed ratio of 2:1, which was sufficient to improve step symmetry. Faster belt speeds that pushed the limit of the patient’s ability, i.e., a slow belt speed determined by the comfortable over ground walking speed (Betschart et al., 2018), instead of the fast belt speed determined by the fastest walking speed (Reisman et al., 2013), was more effective at improving walking speed over ground. Finally, the improvements seen in both studies were retained for at least 1 month, providing encouraging evidence that split-belt walking may be useful for improving walking symmetry over the long term following stroke.

Other forms of motor adaptation have also shown promise for rehabilitation of walking, such as robot applied resistance (Lam et al., 2015), or resistance provided by elastic straps (Blanchette et al., 2012), which have induced adaptations in walking over days. Improvements in over ground walking were seen after patients with spinal cord injury adapted to robot-applied resistances (Lam et al., 2015). Thus, understanding the time course of motor memory induced by such adaptations will be important for planning therapies that produce enduring benefits.

## 5. CONCLUSIONS

Memory of split-belt adaptation was partially retained in all age groups for up to 2 weeks, but less so in adults over 50 years of age. The memory in young children is equal to but not better than young adults. These results suggest that when using split-belt walking in rehabilitation, the interval between therapy sessions to improve walking should be less than 1 week, especially for older patients.

## ACKNOWLEDGEMENTS

This project was supported in part by the Natural Sciences and Engineering Research Council of Canada, grant number 2011-138181 to JFY. Funding to BL was provided by the Women and Children’s Health Research Institute summer research studentship, Alberta Innovates Health Solutions summer studentship, and NSERC Undergraduate Student Research Award over several time periods. We thank Drs Kristin Musselman and Erin Vasudevan for helpful comments on earlier versions of this manuscript. We thank Dr Ming Ye for assistance with statistical modeling and testing, and Dr Jacques Bobet for programing assistance with preparing the figures.

## REFERENCES

Betschart, M., McFadyen, B. J., & Nadeau, S. (2018). Repeated split-belt treadmill walking improved gait ability in individuals with chronic stroke: A pilot study. Physiotherapy Theory and Practice, 34(2), 81–90. https://doi.org/10.1080/09593985.2017.1375055

Betschart, M., McFayden, B. J., & Nadeau, S. (2020). Lower limb joint moments on the fast belt contribute to a reduction of step length asymmetry over ground after split-belt treadmill training in stroke: A pilot study. Physiotherapy Theory and Practice, 36(9), 989–999. https://doi.org/10.1080/09593985.2018.1530708

Blanchette, A., Moffet, H., Roy, J. S., & Bouyer, L. J. (2012). Effects of repeated walking in a perturbing environment: A 4-day locomotor learning study. Journal of Neurophysiology, 108(1), 275–284. https://doi.org/10.1152/jn.01098.2011

Bruijn, S. M., Van Impe, A., Duysens, J., & Swinnen, S. P. (2012). Split-belt walking: Adaptation differences between young and older adults. Journal of Neurophysiology, 108(4), 1149–1157. https://doi.org/10.1152/jn.00018.2012

Day, K. A., Leech, K. A., Roemmich, R. T., & Bastian, A. J. (2018). Accelerating locomotor savings in learning: Compressing four training days to one. Journal of Neurophysiology, 119(6), 2100–2113. https://doi.org/10.1152/jn.00903.2017

Dunbar, M., & Kirton, A. (2018). Perinatal stroke: mechanisms, management, and outcomes of early cerebrovascular brain injury. In The Lancet Child and Adolescent Health (Vol. 2, Issue 9, pp. 666–676). Elsevier B.V. https://doi.org/10.1016/S2352-4642(18)30173-1

Huang, V. S., Haith, A., Mazzoni, P., & Krakauer, J. W. (2011). Rethinking Motor Learning and Savings in Adaptation Paradigms: Model-Free Memory for Successful Actions Combines with Internal Models. Neuron, 70(4), 787–801. https://doi.org/10.1016/j.neuron.2011.04.012

Joiner, W. M., & Smith, M. A. (2008). Long-term retention explained by a model of short-term learning in the adaptive control of reaching. Journal of Neurophysiology, 100(5), 2948–2955. https://doi.org/10.1152/jn.90706.2008

Konczak,, J., Jansen-Osmann, P., Kalveram, K.-T. (2003). Konczak et al 2003 Development of Force Adaptation During Childhood_.pdf (pp. 41–52).

Lam, T, Pauhl, K, Ferguson, A., Krassioukov, A, Eng, J. J. (2015). Training with robot-applied resistance in people with motor-incomplete spinal cord injury: Pilot study. JRRD, Volume 52, Number 1, 52(1), 113–130.

Malone, L. A., & Bastian, A. J. (2010). Thinking about walking: Effects of conscious correction versus distraction on locomotor adaptation. Journal of Neurophysiology, 103(4), 1954–1962. https://doi.org/10.1152/jn.00832.2009

Malone, L. A., & Bastian, A. J. (2016). Age-related forgetting in locomotor adaptation. Neurobiology of Learning and Memory, 128, 1–6. https://doi.org/10.1016/j.nlm.2015.11.003

Malone, L. A., Vasudevan, E. V. L., & Bastian, A. J. (2011). Motor adaptation training for faster relearning. Journal of Neuroscience, 31(42), 15136–15143. https://doi.org/10.1523/JNEUROSCI.1367-11.2011

Mariscal, D. M., Iturralde, P. A., & Torres-Oviedo, G. (2020). Altering attention to split-belt walking increases the generalization of motor memories across walking contexts. Journal of Neurophysiology, 123(5), 1838–1848. https://doi.org/10.1152/jn.00509.2019

Martin, T. A., Keating, J. G., Goodkin, H. P., Bastian, A. J., & Thach, W. T. (1996). Throwing while looking through prisms I. Focal olivocerebellar lesions impair adaptation. Brain, 119(4), 1183–1198. https://doi.org/10.1093/brain/119.4.1183

Mizuno, K., Tsuji, T., Takebayashi, T., Fujiwara, T., Hase, K., & Liu, M. (2011). Prism adaptation therapy enhances rehabilitation of stroke patients with unilateral spatial neglect: A randomized, controlled trial. Neurorehabilitation and Neural Repair, 25(8), 711–720. https://doi.org/10.1177/1545968311407516

Morehead, J. R., Qasim, S. E., Crossley, M. J., & Ivry, R. (2015). Savings upon re-aiming in visuomotor adaptation. Journal of Neuroscience, 35(42), 14386–14396. https://doi.org/10.1523/JNEUROSCI.1046-15.2015

Musselman, K. E., Patrick, S. K., Vasudevan, E. V. L., Bastian, A. J., & Yang, J. F. (2011). Unique characteristics of motor adaptation during walking in young children. Journal of Neurophysiology, 105(5), 2195–2203. https://doi.org/10.1152/jn.01002.2010

Musselman, K. E., Roemmich, R. T., Garrett, B., & Bastian, A. J. (2016). Motor learning in childhood reveals distinct mechanisms for memory retention and re-learning. Learning and Memory, 23(5), 229–237. https://doi.org/10.1101/lm.041004.115

Reisman, D. S., Block, H. J., & Bastian, A. J. (2005). Interlimb coordination during locomotion: What can be adapted and stored? Journal of Neurophysiology, 94(4), 2403–2415. https://doi.org/10.1152/jn.00089.2005

Reisman, D. S., McLean, H., Keller, J., Danks, K. A., & Bastian, A. J. (2013). Repeated split-belt treadmill training improves poststroke step length asymmetry. Neurorehabilitation and Neural Repair, 27(5), 460–468. https://doi.org/10.1177/1545968312474118

Roemmich, R. T., & Bastian, A. J. (2015). Two ways to save a newly learned motor pattern. Journal of Neurophysiology, 113(10), 3519–3530. https://doi.org/10.1152/jn.00965.2014

Torres-Oviedo, G., Vasudevan, E., Malone, L., & Bastian, A. J. (2011). Locomotor adaptation. Progress in Brain Research, 191, 65–74. https://doi.org/10.1016/B978-0-444-53752-2.00013-8

Vasudevan, E. V. L., & Bastian, A. J. (2010). Split-belt treadmill adaptation shows different functional networks for fast and slow human walking. Journal of Neurophysiology, 103(1), 183–191. https://doi.org/10.1152/jn.00501.2009

Vasudevan, E. V. L., Torres-Oviedo, G., Morton, S. M., Yang, J. F., & Bastian, A. J. (2011). Younger is not always better: Development of locomotor adaptation from childhood to adulthood. Journal of Neuroscience, 31(8), 3055–3065. https://doi.org/10.1523/JNEUROSCI.5781-10.2011

Vervoort, D., den Otter, A. R., Buurke, T. J. W., Vuillerme, N., Hortobágyi, T., & Lamoth, C. J. C. (2020). Do gait and muscle activation patterns change at middle-age during split-belt adaptation? Journal of Biomechanics, 99. https://doi.org/10.1016/j.jbiomech.2019.109510

